# Fecal virome-enriched fractions are associated with increased biofilm biomass and distinct bacterial responses in biofilm and planktonic communities in an *in vitro* colon model

**DOI:** 10.64898/2026.09.20.747267

**Authors:** Ania Stenberg Mortensen, Camilla Frost Holm, Henriette Lyng Røder, Torben Sølbeck Rasmussen

## Abstract

The gut microbiota is spatially organized, yet microorganisms at the intestinal mucus surface remain understudied in virome–microbiota research. We investigated how addition of fecal virome-enriched fractions was associated with biofilm biomass and bacterial composition in surface-associated and planktonic communities using a biofilm-adapted *in vitro* colon model, CoMiniGut. Fecal samples were collected from three healthy human donors at two time points approximately one year apart. Communities from both collection time points were fermented for 48 h in reactors containing mucin-coated glass beads. Each community was exposed separately to virome fractions from all three donors, testing autologous and heterologous combinations alongside matched microbiome-only controls. Two parallel CoMiniGut systems provided reactor replicates for each donor–treatment combination. Biofilm biomass was quantified by crystal violet staining, while biofilm-associated and planktonic communities were characterized by 16S rRNA gene sequencing. Across 36 reactor-level observations, biomass was classified as increased in 18, neutral in 11, and decreased in 7. Each observation represented the mean of 5–6 beads normalized to its matched microbiome-only control. Among non-neutral observations, increases were more frequent than decreases (18 versus 7; two-sided exact binomial test, P = 0.043), whereas the magnitude of change did not differ between Increase and Decrease observations (P = 0.220). Virome addition explained a small but significant proportion of bacterial compositional variation (R² = 0.024, P = 0.029) without substantially altering alpha diversity. No interaction between virome treatment and growth mode was detected at the community level, while separate differential-abundance analyses identified different sets of virome-associated taxa in biofilm-associated and planktonic samples. Thus, addition of fecal virome-enriched fractions was associated with a directional bias toward increased biofilm biomass, although the responsible retained components were not identified.

## 1. Introduction

The human gut microbiota is spatially organized, with differences in microbial composition and activity between the intestinal lumen and the mucus-covered epithelium [1,2]. The gut also contains a diverse viral community, referred to as the gut virome. It is dominated by bacteriophages, hereafter referred to as phages, but also includes viruses infecting eukaryotic and archaeal hosts. Phages can alter bacterial abundance, function, and community composition through infection, selection, and genetic exchange, with indirect effects on other community members [3–6]. However, gut virome–microbiota studies commonly rely on fecal samples [7], and comparatively few have examined the mucosal–luminal interface or characterized biofilm-associated bacterial and viral communities separately [7,8].

The intestinal epithelium is covered by mucus, which limits contact with luminal microorganisms. In the colon, mucus consists of a dense inner layer that restricts bacterial access to the epithelium and a looser outer layer that provides nutrients and attachment sites for microorganisms [9,10]. Bacterial communities associated with the intestinal mucosa can consequently differ from those detected in feces or the lumen [11]. In the murine intestine, spatial analysis identified a mucus-embedded, biofilm-like bacterial community that was compositionally distinct from the adjacent luminal community [12]. Mucus can also affect phage–bacterium interactions. Retention of phages near the mucus surface may increase infection and killing of susceptible bacteria and thereby limit bacterial access to the epithelium [13]. Mucus adherence has increased phage persistence and antimicrobial activity in fish and murine models, although mucin can also alter bacterial attachment, metabolism, and phage susceptibility in a system-dependent manner [14–16].

Bacteria at the mucus surface can form biofilm-like communities with properties that differ from planktonic populations [17,18]. Their extracellular matrix and local differences in bacterial growth can restrict phage movement, reduce access to susceptible bacteria, and alter the spread of infection [19–21]. In the intestine, this matrix is integrated with host-derived mucus. We recently suggested a new term “*mucofilm*” that describes this combined environment of mucus, microorganisms, bacterial matrix, and phages [22]. Models combining mucin with surface-attached bacterial growth may therefore reveal phage-bacterium interactions that depend on spatial structure and biofilm formation and are not normally observed in purely planktonic systems.

Phages may increase or decrease biofilm biomass through several mechanisms. Lysis can release cellular material, including extracellular DNA, that may become incorporated into the biofilm matrix [23,24]. Phage exposure can also alter bacterial physiology or select for bacteria with increased matrix production and altered phage susceptibility [25,26]. In mixed communities, changes in susceptible bacteria can affect species outside the direct phage host range through altered bacterial interactions [5,27]. Complex viromes may therefore affect biofilm biomass through both direct infection and indirect community changes. In the gut, phages occur as part of complex viral communities rather than as individual isolates. Consequently, the fecal virome contains a diverse mixture of phages that may influence several bacterial populations simultaneously. Interest in these community-level effects has developed partly through fecal microbiota transplantation (FMT), which transfers bacteria, phages, metabolites, and other donor-derived components. Donor phages can persist in recipients, and their transfer has been associated with changes in bacterial and viral communities and, in some studies, treatment outcome [28–32]. Fecal filtrate transfer can also produce measurable effects without transferring intact donor bacteria. However, these preparations retain phages together with metabolites, proteins, bacterial DNA, and other components, preventing the effects from being attributed specifically to phages [33,34]. Fecal virome transplantation (FVT) uses additional processing steps to enrich virus-like particles and reduce smaller soluble metabolite components [33,35–39]. Animal studies have shown that fecal virome addition can alter established bacterial communities and, in some models, host phenotypes [34,37,40,41]. Because gut bacterial and viral communities differ substantially among individuals, responses may depend on the specific bacterial community-virome pairing [42,43]. Autologous pairings contain a bacterial community and virome from the same donor, whereas heterologous pairings combine a bacterial community and virome from different donors. While fecal virome studies have demonstrated community-level effects, it remains largely unknown whether biofilm-associated and planktonic bacterial populations respond similarly to virome interventions.

Here, we investigated whether addition of fecal virome-enriched fractions was associated with changes in biofilm biomass and bacterial composition using a biofilm-adapted version of an *in vitro* colon model called CoMiniGut [44]. Mucin-coated porous glass beads enabled separate characterization of biofilm-associated and planktonic communities from the same reactors. Bacterial communities and virome fractions were obtained from three healthy human donors at two sampling time points one year apart. Each bacterial community was exposed to virome fractions from all three donors, testing autologous and heterologous combinations alongside matched controls without virome addition. We hypothesized that addition of the fecal virome-enriched fractions would alter biofilm biomass and bacterial composition and that different bacterial taxa would respond in biofilm-associated and planktonic communities. Because most virome studies focus on fecal communities, this approach enabled us to assess whether virome-mediated responses differ between spatial niches within the same microbial ecosystem.

## 2. Material and methods

### 2.1 Experimental design

Fecal samples were collected from three healthy human donors at two time points approximately one year apart (Year 1, 2024; Year 2, 2025) providing a single sample pr. Year pr. donor, and a virome-enriched fraction was extracted from each sample, yielding Virome 1, Virome 2, and Virome 3 from Donors 1–3, respectively. Because the fecal inocula naturally contained resident phages, the experimental intervention was the addition of a concentrated virome fraction from either the same donor or a different donor. Fecal microbiota were fermented in the *in vitro* CoMiniGut system [44] under five treatment conditions: supplementation with Virome 1, Virome 2, or Virome 3; a microbiome-only control containing fecal inoculum without additional virome; and a media-only control without fecal inoculum. Two CoMiniGut systems were run in parallel, providing two reactor replicates per donor–treatment combination within each experimental year. Planktonic samples (free floating bacteria) were collected at 0 h, 21 h, and 48 h. The virome-enriched fraction was added after 20 h of fermentation to allow initial colonization and early biofilm establishment on the mucin-coated beads before intervention. Previous studies have demonstrated rapid colonization of mucin surfaces by complex gut microbial communities and subsequent development of biofilm-like communities, while biofilm formation and maturation continue over time [45]. The remaining 28 h of fermentation allowed the supplemented virome fraction to interact with and potentially influence ongoing biofilm development and bacterial community composition

### 2.2 Fecal samples and virome preparation

All stool samples were collected from three healthy in-house donors (D1–D3; Table 1) with written consent in accordance with ethical committee (The Committees on Health Research Ethics in the Capital Region of Denmark) approval H-20028549. Donors reported no chronic disease, antibiotic or antiviral use within three months prior to sampling, or use of any prescribed medication. Approximately 25 g of fresh stool was homogenized in 25 mL glycerol:NaCl solution, filtered through a BagPage filter bag (280 micrometer filter porosity, InterScience, Ref 121 025), and aliquoted (1800 µL) into cryotubes. Fecal suspensions were prepared by homogenization in glycerol:NaCl solution. The required volume varied between donors due to differences in fecal consistency and viscosity. To ensure viability of the strict anaerobic bacteria, fecal samples were brought to an anaerobic work bench maximum 10 minutes after sampling (Don Whitley A85, atmosphere 80% H_2_, 10% CO_2_, 10% N_2_), and all solutions were prepared anoxically by boiling and flushing with 100% N_2_ for 20 min. Samples were stored at −80 °C. For virome preparation, the fecal suspension was thawed, and subsequently 1 mL (1000 µL) of fecal suspension was diluted in SM buffer to a final volume of 5 mL and centrifuged twice at 4,402 × g for 30 min at 4 °C. After each centrifugation, the supernatant was transferred to a new sterile tube without disturbing the pellet. The resulting supernatant was passed through a 0.45 µm PES filter to remove most bacterial cells and larger particles and was subsequently concentrated using a Pierce protein concentrator PES with a 100 kDa molecular-weight cut-off at 2,500 × g to a final volume of approximately 1 mL [35,38]. This procedure was used to enrich virus-like particles and reduce smaller soluble components. Concentrated fractions were stored at 4 °C for up to one week before use. Viral taxonomical composition, virus-like particle concentration, and infectivity were not determined for the individual fractions.

**Table 1.** Donor characteristics and sample metadata. Samples were collected from three healthy donors at two sampling time points approximately one year apart. Donors reported no chronic disease, antibiotic use within three months prior to sampling, or use of prescribed medication. Age, Bristol Stool Scale (BSS) scores are shown for each sampling time point. Fecal samples were processed as described in the Materials and Methods section.

| **Donor** | **Season 1** | **Season 2** |
|---|---|---|
| **Donor 1 (D1)** | 37 y/o male<br>Bristol Stool Scale: 4 | 38 y/o male<br>Bristol Stool Scale: 4 |
| **Donor 2 (D2)** | 31 y/o male<br>Bristol Stool Scale: 4 | 32 y/o male<br>Bristol Stool Scale: 5 |
| **Donor 3 (D3)** | 24 y/o female<br>Bristol Stool Scale: 5 | 25 y/o female<br>Bristol Stool Scale: 3 |

### 2.3 In vitro colon model (CoMiniGut)

Anaerobic BCM (Basic Colon Media) media was adapted from basal media by Wiese et al. [44]. The media was prepared by dissolving bile salts (0.5 g/L), NaHCO (2 g/L), MES hydrate (3.94 g/L), Tween 80 (2 mL/L), M-SHIME nutritional media (14.6 g/L), resazurin (1 mL/L of 1% w/v stock), and appropriate volumes of salt stock solutions (NaCl, K HPO , KH PO , MgSO ·7H O, CaCl ·2H O, NaHCO ; each prepared at 10 g/L or as noted) in Milli-Q water. The pH was adjusted to ∼5.7 during BCM preparation and the volume brought to 1 L. The media was rendered anaerobic by boiling and flushing with 100% N for ≥15 min using an Anaerobic Gassing Unit, then autoclaved at 121 °C for 20 min. Following cooling, hemin (2.0 mL/250 mL) and vitamin K (400 µL/250 mL) stock solutions were added. BCM media was stored at 4 °C in the dark for up to one month. Prior to fermentation, the medium was adjusted to pH 6.6–6.8. Fermentation was conducted using the *in vitro* CoMiniGut system as described by Wiese et al. [44]. The setup was adapted to support surface-associated growth by extending fermentation from 24 to 48 h and incorporating mucin-coated surfaces. Because biofilm biomass changes dynamically during development and dispersal, the 48-h endpoint should be interpreted as a defined sampling time rather than as a measure of maximal biofilm formation [46,47]. Each reaction vial was loaded with one magnetic stirrer and 11 mucin-coated glass beads, sealed under anaerobic conditions with an anaerobic sachet, and filled with 9.5 mL (10 mL in media only controls) BCM media and 0.5 mL fecal suspension (1:5 dilution in PBS) to a final working volume of 10 mL. Fermentation was run for 48 h at 37 °C with automated pH maintenance at 6.6–6.8 via controlled NaOH addition, controlled through custom MATLAB scripts (ver. R2015a; The MathWorks, Inc., Natick, MA, USA). Concentrated virome (500 µL) was added at t = 20 h in accordance with experimental setup.

### 2.4 Biofilm formation and sampling on mucin-coated glass beads

Porcine gastric mucin powder (Sigma-Aldrich) was sterilized by incubation in 96% ethanol at 65 °C for 24 h, followed by ethanol evaporation and reconstitution at 1% (w/v) in sterile demineralized water. Borosilicate glass beads (4 mm; ROBU) were immersed in the mucin solution for 1 h at 120 rpm, after which excess mucin was removed and beads were dried under sterile conditions. Planktonic samples (1800 µL) were collected at t_0_, t_21_ and t_48_ using a syringe needle and luerlock syringe, aliquoted into cryotubes, and stored at −60 °C. At termination, five beads per reactor were recovered with sterile forceps, transferred to cryotubes containing planktonic material from the corresponding reactor, and stored at −60°C until processing for DNA extraction. Before DNA extraction, the beads were removed from the storage suspension and washed as described below. The six remaining beads in each reactor were transferred individually to a sterile 96-well plate for crystal violet assay. Biofilm formation on mucin-coated glass beads was quantified by crystal violet assay. Beads were washed three times with SM buffer (180 µL) to remove non-adherent bacteria, stained with 1% crystal violet (180 µL) for 45 min at room temperature, and washed with Milli-Q water until the water ran clear. Stain was solubilized in 96 % ethanol (180 µL) for 1 h, and 100 µL of the eluate was transferred to a fresh 96-well plate for absorbance measurement at 590 nm (Varioskan LUX).

### 2.5 DNA extraction and 16S rRNA gene sequencing

Bacterial DNA was extracted from original fecal samples, planktonic CoMiniGut samples, and bead-associated biofilm samples using the Bead-Beat Micro AX Gravity kit (cat. no. 106-100-mod1; A&A Biotechnology), with sample-specific preparation as described below. For planktonic and original fecal samples, 1 mL of sample was centrifuged at 10,000 × g for 10 min. The resulting pellet was resuspended in PBS to a final volume of 80–100 µL before DNA extraction. For bead-associated biofilm samples, the five mucin-coated beads were transferred from the storage suspension to sterile 2 mL tubes using sterile forceps and washed three times with SM buffer to remove planktonic cells. Bead-associated material was released by covering the beads with 0.1% (w/v) dithiothreitol in PBS, followed by vortexing for 1 min, shaking at 300 rpm for 1 min, and vortexing for an additional 1 min. The beads were then removed using sterile forceps. The released material was centrifuged at 10,000 × g for 10 min, and the resulting pellet was resuspended in PBS to a final volume of 80–100 µL before DNA extraction. The prepared samples were processed using the Bead-Beat Micro AX Gravity kit according to the manufacturer’s instructions. Samples were incubated with lysozyme and mutanolysin, followed by proteinase K treatment and mechanical disruption by bead beating for 60 s at 4.5 m/s. DNA from planktonic and original fecal samples was eluted in 60 µL elution buffer, whereas DNA from bead-associated samples was eluted in 40 µL. DNA extracted from bead-associated samples was purified using magnetic binding beads to remove residual dithiothreitol before 16S rRNA gene amplification. Media-only control samples were processed alongside biological samples throughout DNA extraction and sequencing. For each extraction batch (22+2), one negative extraction control and one mock community control were included. Negative extraction controls consistently yielded fewer than 1000 reads and were removed during bioinformatic filtering. Mock community controls were used to verify sequencing performance and negative extraction controls were included for quality-control purposes. The final purified DNA was stored at -60 °C and the DNA concentration was determined using the Qubit HS Assay Kit on the Qubit 4 Fluorometric Quantification device (Invitrogen). Near-full-length 16S rRNA gene sequencing and initial sequence processing were performed as previously described [48], with the sample-specific procedures and quality thresholds detailed below. In brief, the 16S rRNA gene was amplified by polymerase chain reaction (PCR) with primers targeting conserved regions flanking the hypervariable regions V1-V8. The initial PCR (PCR1) reaction mixture included PCRBIO HiFi polymerase and PCRBIO buffer (PCR Biosystems Ltd.), primer mix, genomic DNA, and nuclease-free water. Gel electrophoresis was used to verify the size of the PCR products that subsequently were barcoded by an additional PCR (PCR2) reaction using the same reagents but with barcoded primers. The final PCR products were purified using AMPure XP beads (Beckman Coulter) and pooled in equimolar concentrations. The pooled barcoded amplicons were ligated according to 1D genomic DNA using a ligation protocol (SQK-LSK109) to complete library preparation for sequencing on a R10.4.1 flow cell. Data generated by the MinION were collected using MinKnow software v19.06.8. The Guppy v3.2.2 basecalling toolkit was used to base call raw fast5 to fastq. Porechop v0.2.2 was used for adapter trimming and sample demultiplexing (https://github.com/rrwick/Porechop). Sequences containing quality scores (fastq files) were quality corrected using NanoFilt (q ≥ 10; read length > 1Kb). Taxonomy assignment of quality corrected reads against Greengenes (13.8) database was conducted using uclast method implemented in parallel_assign_taxonomy_uclust.py (QIIME v1.9.1) [48].

### 2.6 Bioinformatics and statistical analysis

Individual bead measurements were treated as technical replicates, and the mean OD590 of 5-6 beads was used to estimate biofilm biomass for each reactor (biological replicate). Media-only reactors were included as process controls to monitor background biomass formation within the CoMiniGut system but were not used for normalization of crystal violet measurements. Instead, virome-treated reactors were compared with the matched microbiota-only control.

Biofilm biomass was analyzed as a continuous outcome using a linear mixed effects model on the raw OD590 values, with donor and virome treatment as fixed effects and CoMiniGut system and experimental year as random effects (OD ∼ Donor + Treatment + (1 | Machine) + (1 | Year)). This model was compared with a model including a donor by treatment interaction, and the contribution of each random effect was assessed by removing it from the full model. Model assumptions were evaluated visually using residual versus fitted plots, Q-Q plots, and random effect distributions, with no major violations observed. Statistical significance was assessed at α = 0.05. To describe the direction of the virome response, for each virome treatment, fold change was calculated by dividing its mean OD_590_ by that of the corresponding microbiome-only control matched by fecal donor, experimental year, and CoMiniGut system. The matched microbiome-only control therefore corresponded to a fold change of 1.0. Observations were descriptively classified as Increase if the mean fold change minus its standard error exceeded 1.0, Decrease if the mean fold change plus its standard error was below 1.0, and Neutral if the standard-error interval encompassed 1.0. These categories were used descriptively and do not represent statistical significance tests for individual reactor replicates. The frequencies of Increase and Decrease observations were compared using a two-sided exact binomial test, excluding Neutral observations. For observations classified as Increase or Decrease, the magnitude of change was calculated as the absolute log2 fold change, which is symmetric around the microbiome-only reference value (log2 fold change = 0), and compared between groups using a two-sided Wilcoxon rank-sum test. All analyses were performed in R version 4.3.1 and visualized using ggplot2. Bacterial community composition was assessed by PERMANOVA using adonis2 in the vegan package on Bray–Curtis dissimilarities, with within-group dispersion evaluated by PERMDISP. Read counts were normalized by cumulative sum scaling prior to beta-diversity analysis. Alpha diversity, measured as observed OTU richness and Shannon diversity, was calculated after normalization to the median sequencing depth. Group comparisons were performed using pairwise Wilcoxon rank-sum tests with false-discovery-rate correction. Differential abundance was assessed using DESeq2 with donor identity and virome treatment included in the design (∼ Donor + Virome). P-values were adjusted using the Benjamini– Hochberg false discovery rate, and OTUs were considered differentially abundant at adjusted P < 0.05 and absolute log2 fold change >0.5. Analyses were performed in R and visualized using ggplot2.

## 3. Results

### 3.1 Biofilm formation in the in vitro CoMiniGut system

Crystal violet staining detected surface-associated biomass on the mucin-coated glass beads across the donor-derived fermentations. The effect of adding the fecal virome-enriched fractions was evaluated by calculating fold change in mean OD590 for each treated reactor relative to its matched microbiome-only control (Figure 2A). Across the 36 reactor replicates, biofilm biomass was classified as increased in 18, neutral in 11, and decreased in 7. Among the 25 observations classified as either Increase or Decrease, increases were more frequent than decreases (18 versus 7; two-sided exact binomial test, P = 0.043). However, the absolute deviation of the fold change from the microbiome-only reference value of 1.0 did not differ significantly between Increase and Decrease observations (two-sided Wilcoxon rank-sum test, P = 0.220; Figure 2B). These results indicate a directional bias towards increased biofilm biomass rather than a difference in the magnitude of the relative change. Decreases were primarily observed for donor 3 in year 2, highlighting variation across donors and experimental rounds.

**Figure 1.**
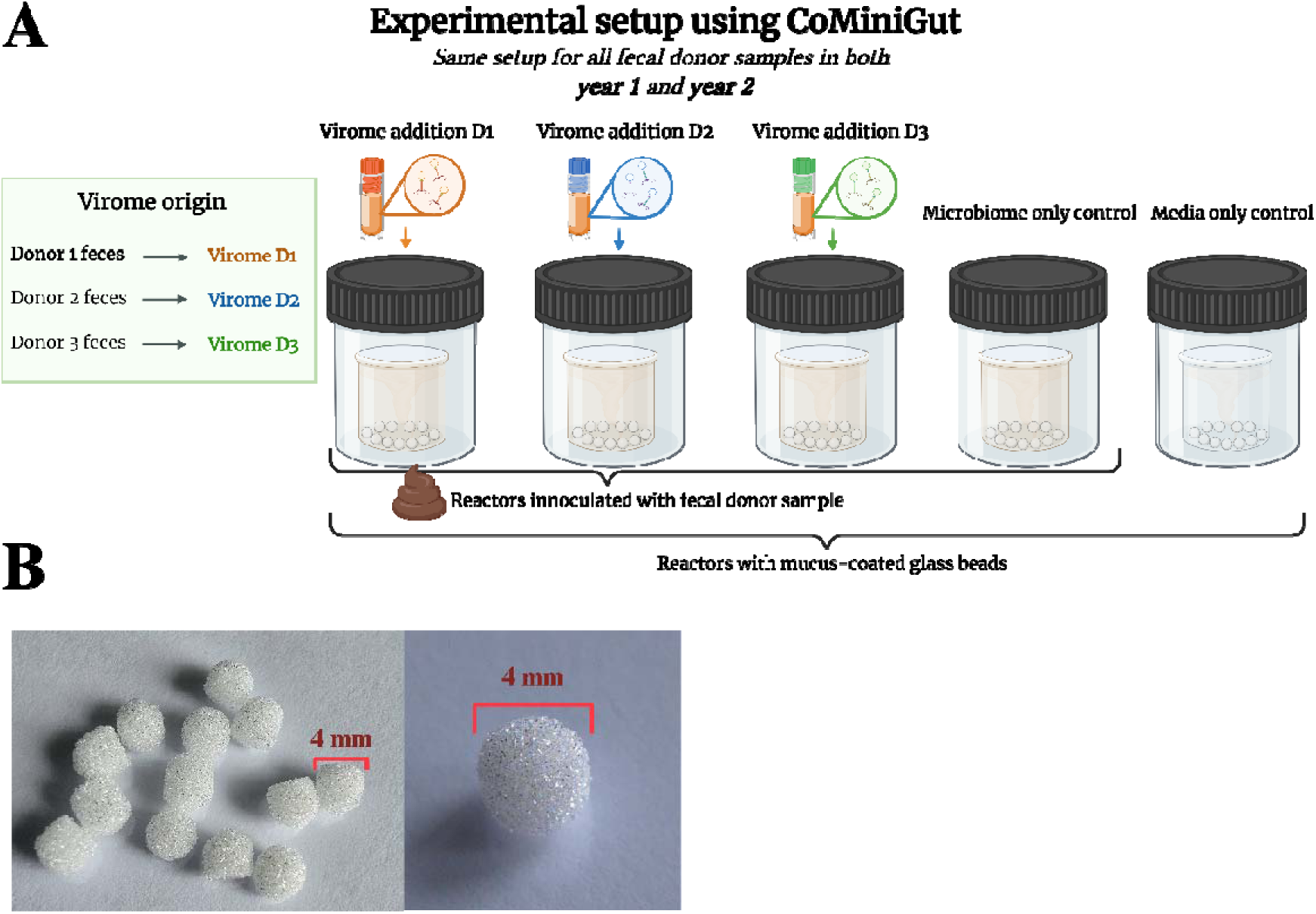
Experimental design of the biofilm-adapted CoMiniGut in vitro colon model. A) Fecal virome-enriched fractions (Explained in section 2.2) were prepared from three healthy human donors (Donor 1-3), yielding Virome 1, Virome 2, and Virome 3. For each donor, parallel CoMiniGut reactors containing mucin-coated glass beads were inoculated with fecal material and supplemented with Virome 1, Virome 2, Virome 3, or microbiome-only (control). An additional reactor containing medium and mucin-coated beads only served as a negative control. Two CoMiniGut systems were run in parallel, generating two reactor replicates for each donor-treatment combination. The entire experiment was repeated at two sampling time points approximately one year apart. B) ROBU Vitrapor porous borosilicate glass beads 4 mm used to create a mucin surface for these experiments. Left: multiple beads. Right: Close-up of one bead to show its porous surface. Created in BioRender. Rasmussen, T. S. (2027) https://BioRender.com/eill64b

**Figure 2.**
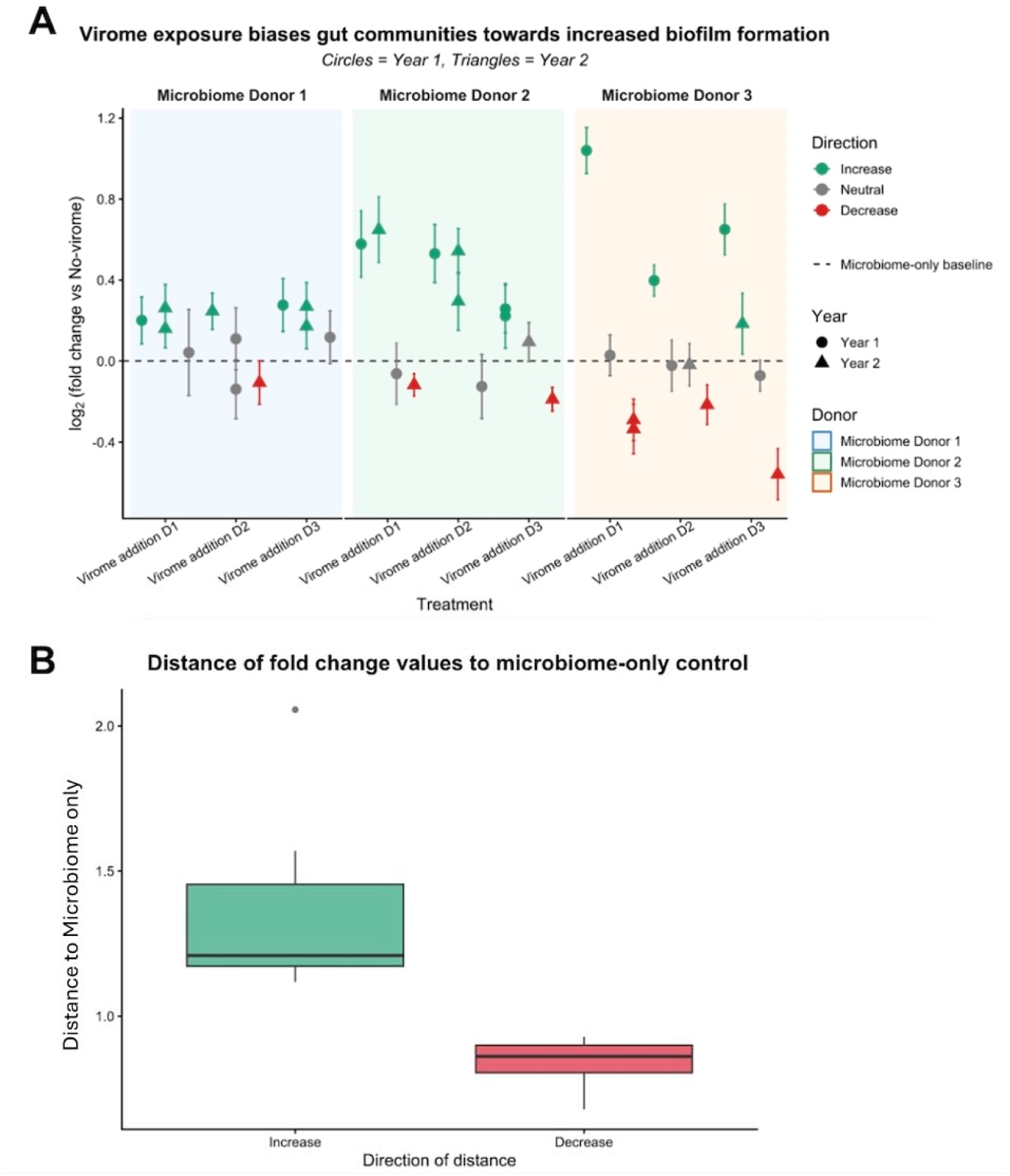
Changes in biofilm biomass associated with addition of fecal virome-enriched fractions in the CoMiniGut system. (A) Fold change in crystal violet-stained biofilm biomass following addition of fecal virome-enriched fractions relative to the corresponding microbiome-only control. Each point represents one reactor replicate and the mean OD590 measured across 5–6 mucin-coated glass beads, normalized to the microbiome-only control matched by fecal donor, collection time point, and CoMiniGut system. The dashed horizontal line at 1.0 represents the matched microbiome-only reference. Observations were descriptively classified as Increase, Decrease, or Neutral based on whether the standard-error interval was above, below, or encompassed 1.0, respectively. (B) Absolute deviation of the fold change from 1.0 for observations classified as Increase or Decrease; Neutral observations were excluded. The groups did not differ significantly in magnitude (two-sided Wilcoxon rank-sum test, P = 0.220). Media-only control values were not subtracted before calculation of fold changes.

### 3.2 Microbiome shifts in planktonic and biofilm-associated samples

Bacterial community composition in CoMiniGut-fermented samples was strongly associated with donor identity and fecal collection time point, which explained 36% and 10% of the total compositional variation, respectively (Supplementary Table 1). Multivariate dispersion differed among donors but not between collection time points. The donor-associated PERMANOVA effect may therefore reflect differences in both average community composition and within-donor variability.

Fermented communities differed significantly from the original donor fecal communities (R² = 0.11, P < 0.001; Figure 3A and Supplementary Table 1). Nevertheless, the fermented communities remained more compositionally similar to their respective source fecal communities than to the CoMiniGut background microbiota (Supplementary Figure 1). Donor-associated community characteristics therefore remained detectable after 48 h of fermentation, despite the compositional changes associated with the CoMiniGut conditions.

**Figure 3.**
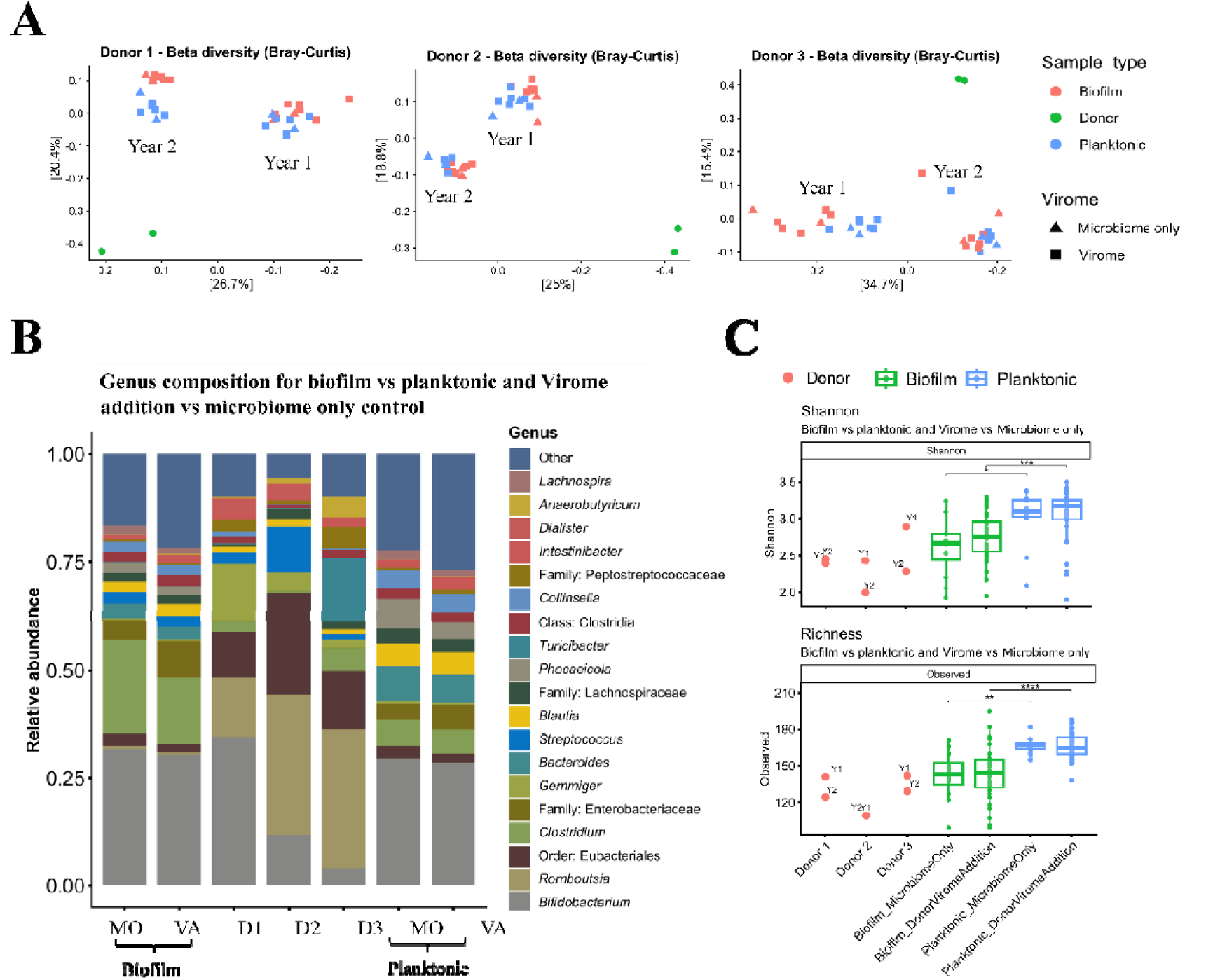
Bacterial community composition and diversity of CoMiniGut fermented samples. **(A)** Principal Coordinates Analysis (PCoA) plots based on Bray-Curtis dissimilarity for all CoMiniGut fermentations including fecal suspension, divided by donor identity: Donor 1 (A), Donor 2 (B), and Donor 3 (C). Shape indicates virome treatment (Microbiome-only: triangle, Virome: square) and color indicates sample type (Biofilm: red, Planktonic: blue, Donor: green). **(B**) Stacked bar plot showing genus-level relative abundance (taxa with mean relative abundance ≥1%) for biofilm and planktonic samples under Microbiome-only (MO) and Virome addition (VA) treatments, alongside original donor fecal samples (D1, D2, D3). **(C)** Alpha diversity of donor fecal samples, biofilm, and planktonic communities under Microbiome-only and Virome addition conditions, shown as observed OTU richness and Shannon index. Planktonic samples showed significantly higher diversity than biofilm for both richness (mean 167 vs 143, **P<0.01) and Shannon index (mean 3.0 vs 2.7, *P<0.05). Statistical comparisons by two-sided Wilcoxon rank-sum test with FDR correction (*P<0.05, **P<0.01, ***P<0.001, ****P<0.0001).

Biofilm-associated and planktonic samples differed significantly in both alpha diversity and bacterial community composition. Planktonic samples had higher Shannon diversity than biofilm-associated samples under both microbiome-only and virome-addition conditions (FDR-adjusted P < 0.05 and P < 0.001, respectively; Figure 3C). Observed OTU richness was likewise higher in planktonic samples under microbiome-only and virome-addition conditions (FDR-adjusted P < 0.01 and P < 0.0001, respectively; Figure 3C). Bacterial community composition also differed significantly between the two sample types (PERMANOVA, R² = 0.069, P < 0.001; Supplementary Table S1). Multivariate dispersion also differed significantly between biofilm-associated and planktonic samples (PERMDISP, P < 0.001; Supplementary Table S1). The PERMANOVA result for sample type may therefore reflect differences in both average community composition and within-group variability.

The taxonomic profiles of both sample types were dominated by *Bifidobacterium*. Among the remaining taxa displayed in Figure 3B, *Clostridium* and members of the family *Enterobacteriaceae* accounted for larger proportions of the biofilm-associated profiles, whereas *Bacteroides* and *Blautia* accounted for larger proportions of the planktonic profiles. *Streptococcus* was visible in the biofilm-associated but not the planktonic profiles shown in Figure 3B.

Addition of the fecal virome-enriched fractions was associated with a small but statistically significant difference in bacterial community composition between virome-addition and microbiome-only samples (PERMANOVA, R² = 0.024, P = 0.029; Supplementary Table S1). No interaction between virome treatment and sample type was detected at the overall community level (R² = 0.0089, P = 0.99; Supplementary Table S1). Multivariate dispersion did not differ between virome-addition and microbiome-only samples (PERMDISP, P = 0.76; Supplementary Table S1), indicating that the virome-associated PERMANOVA result was unlikely to be explained by unequal within-group dispersion.

Separate differential-abundance analyses identified distinct sets of bacterial OTUs associated with addition of the fecal virome-enriched fractions in planktonic and biofilm-associated samples (Supplementary Figure S2). The identified OTUs were generally of low relative abundance, supporting a subtle taxonomic response rather than extensive community restructuring.In planktonic samples, addition of the fecal virome-enriched fractions was associated with increased relative abundances of *Alistipes* (virome mean, 2.63%; microbiome-only, 0.98%) and *Coprococcus* (2.74% versus 0.54%), whereas *Phocaeicola* was less abundant (0.75% versus 1.47%). In biofilm-associated samples, *Coprococcus* was also more abundant following virome addition (1.79% versus 0.53%), while an OTU assigned to the order *Eubacteriales* was less abundant (1.93% versus 2.80%). The magnitude of these descriptive differences varied among the three virome fractions, with particularly pronounced changes under Virome 2 for some taxa.

## 4. Discussion

Fecal virome addition shifted biofilm biomass towards increased values in the *in vitro* CoMiniGut model. Across 36 reactor replicates, biomass was classified as increased in 18, neutral in 11, and decreased in 7. Increases were more frequent than decreases among non-neutral observations, whereas the magnitude of the relative change did not differ between Increase and Decrease observations. Addition of the fecal virome-enriched fractions was also associated with a small but significant change in bacterial community composition without a substantial change in alpha diversity. The overall change in composition was similar in biofilm and planktonic communities, but different bacterial taxa were affected in the two communities. Addition of the fecal virome-enriched fractions was therefore associated with changes in both biofilm biomass and bacterial composition, although the present design does not establish which retained component of the fractions caused these changes. Although absolute biofilm levels were modest, surface-associated biomass was consistently detected across all donor communities, indicating that the mucin-coated bead system reproducibly supported biofilm formation under the experimental conditions.

Although phages are commonly associated with bacterial killing and biofilm removal, they can either reduce or increase biofilm biomass depending on the phage, bacterial host, biofilm stage, and surrounding bacterial community [21,49], providing one biologically plausible explanation for the observed biomass pattern. The preparation workflow was developed to enrich virus-like particles and has previously been shown to recover infective phages from fecal material, although infectivity was not assessed in the present study and recovery depends on the phage, sample matrix, and ultrafilter used [35,38]. Lysis can release cellular material, including extracellular DNA, that may become incorporated into the biofilm matrix. Consistent with this mechanism, spontaneous prophage induction promoted biofilm formation in an *Actinomyces odontolyticus* model through the release of host-derived extracellular DNA [23]. Low phage exposure can also change bacterial physiology and stimulate biofilm formation, while longer exposure can select for bacteria with increased matrix production and altered phage susceptibility [25,26]. In mixed bacterial communities, infection of susceptible bacteria can also affect bacteria outside the direct phage host range by changing competition and other bacterial interactions [5,50]. Despite the potential contribution of multiple mechanisms, the directional bias towards increased biomass observed across donors and experimental conditions suggests that fecal virome addition can influence the organization of surface-associated gut microbial communities. The crystal violet results should be interpreted as changes in total material retained on the beads, not as direct evidence of changes in viable bacterial numbers. Crystal violet stains bacterial cells and extracellular material and does not distinguish between viable cells, dead cells, and biofilm matrix [51,52]. Phage-mediated lysis could therefore increase the amount of extracellular material on the beads without increasing bacterial cell numbers. Changes in bacterial attachment or matrix production could produce a similar result. Future studies should combine crystal violet staining with viable-cell counts, absolute bacterial quantification, measurements of extracellular DNA and polysaccharides, and microscopy of the beads.

Virome addition explained only a small part of the variation in bacterial composition. Bacterial donor, biofilm versus planktonic growth, and experimental round explained more of the variation. This is not surprising because gut bacterial communities retain strong donor-specific features [42,53,54], and CoMiniGut reproduces donor-dependent fermentation responses with good experimental reproducibility [44,55]. Virome addition therefore changed communities that were already strongly shaped by their bacterial donor and the experimental conditions. The lack of a substantial change in alpha diversity by virome addition further suggests that the viromes mainly changed the relative distribution of existing bacteria rather than causing broad loss of bacterial diversity. Previous fecal virome studies have also found changes in established bacterial communities and specific bacterial taxa without complete restructuring of the recipient community [37,40,41].

Although virome addition explained a similar proportion of variation in biofilm-associated and planktonic communities, the bacterial taxa contributing to this response differed. This suggests that bacterial growth mode influences how communities respond to fecal virome addition. In other words, similar community-level effects can arise through different taxonomic pathways in different ecological niches. Bacteria attached to the mucin-coated beads and bacteria growing in suspension experience different conditions and can differ in growth, metabolism, surface structures, and phage susceptibility. The biofilm matrix can restrict phage movement and access to susceptible bacteria, while differences in bacterial growth within the biofilm can change the spread of infection [19–21,56]. These differences may explain why virome addition affected different taxa in biofilm and planktonic communities.

The present results extend this concept by showing that even when the same bacterial community and virome are present within the same reactor, bacterial responses can differ between biofilm-associated and planktonic compartments. Gut virome research has mainly used fecal samples, although viruses associated with the intestinal mucosa can differ from those detected in stool. Studies of the colonic mucosal–luminal interface identified biofilm-associated phages that were not detected by fecal sampling [7,8]. The present study tested a different question by adding the same fecal virome fractions to reactors containing both a mucin-coated surface and a planktonic community. The different taxonomic responses indicate that analysis of the planktonic phase alone may miss bacterial changes associated with biofilm growth.

The mucin-coated beads provide a simplified model of the intestinal mucus surface. They do not reproduce the complete mucus layer, which is continuously produced and renewed and contains host proteins, antimicrobial molecules, immune factors, and chemical gradients [57,58]. The model also lacks epithelial and immune cells, intestinal flow, and host-mediated clearance. The biofilm-associated communities should therefore be viewed as biofilms growing on a mucin-coated surface, not as a complete reconstruction of the intestinal mucosa. The main advantage is that biofilm and planktonic communities develop in the same reactor and are exposed to the same inoculum, virome fraction, medium, and fermentation conditions. This allows the two communities to be compared directly.

Testing all bacterial community–virome combinations from three donors reduced the risk that the overall biomass pattern was caused by a single donor combination. However, the study was not designed to establish a general difference between autologous and heterologous combinations. More donors are needed to identify bacterial or viral features that predict increased or decreased biofilm biomass. The two sampling time points also add variation within donors but do not represent independent donor groups. Bacterial and viral gut communities can change over time [43,59]. Technical conditions may also have differed between experimental runs. The effect of experimental round may therefore reflect both changes in donor material and technical variation.

Fecal virome-enriched fractions have previously been shown to alter bacterial communities in both *in vivo* [60] and defined *in vitro* systems [61]. However, viral composition, particle concentration, and infectivity were not determined in the individual fractions used here. Also, ultrafiltration enriches material according to physical retention rather than viral identity.

Although smaller soluble components were reduced during processing, larger proteins, extracellular vesicles, or other retained fecal components may have remained. Consequently, the observed effects should be interpreted as responses to fecal virome-enriched fractions rather than direct evidence of active phage infection. While infectivity assays can confirm the presence of phages capable of infecting selected bacterial hosts, such assays capture only a subset of interactions occurring within complex microbial communities [62]. Likewise, individual plaque-forming phages do not necessarily represent those that contribute most strongly to community-level dynamics in complex virome-host systems [63]. Future studies combining viral characterization, infectivity assessment, and appropriate controls such as ultrafiltration permeates or selectively virus-inactivated fractions will be important for establishing the relative contribution of viral and non-viral components to the observed community responses.

The 16S rRNA gene amplicon data provide relative bacterial composition but do not measure absolute bacterial abundance or strain-level changes. DNA yields from bead-associated samples were substantially lower than those from planktonic samples, and differences in sample processing and DNA recovery may therefore have contributed to the observed differences in diversity and composition between the two sample types. An increase in the relative abundance of one taxon may reflect its own growth, a decrease in another taxon, or both [64,65]. This is particularly important when comparing the sequencing results with the crystal violet measurements. Crystal violet quantified total material on the beads, whereas sequencing measured the relative composition of bacterial DNA. Increased biomass could therefore result from more bacterial cells, increased matrix production, or material released through lysis without requiring a large change in bacterial composition. Future studies should combine crystal violet measurements with absolute and viable bacterial quantification, strain-resolved community analysis, and direct measurements of biofilm matrix components.

## 5. Conclusion

Addition of fecal virome-enriched fractions shifted biofilm biomass towards increased values and was associated with a small but significant difference in bacterial community composition in a biofilm-adapted *in vitro* colon model. Although the overall compositional shift was comparable between biofilm-associated and planktonic communities, different bacterial taxa contributed to the response in the two growth forms. These findings show that addition of fecal virome-enriched fractions is associated with changes in bacterial community organization during surface-associated and planktonic growth. The results support the inclusion of mucus-relevant, surface-associated compartments in experimental studies of gut virome-microbiota interactions and identify biofilm biomass as an important outcome when evaluating virome-based interventions.

## Data availability

Raw 16S rRNA gene sequencing reads can be found at European Nucleotide Archive under ID PRJEB128576. Scripts and data sheets used to generate the results can be accessed through Mendeley Data (doi: 10.17632/wvhx6hm4xd.1).

## CRediT authorship contribution statement

**Ania Stenberg Mortensen:** Conceptualization, Methodology, Investigation, Writing – original draft, Writing – review and editing. **Camilla Frost Holm:** Methodology, Investigation, Formal analysis, Visualization. **Henriette Lyng Røder:** Conceptualization, Methodology, Supervision, Project administration, Writing – review and editing, Funding acquisition. **Torben Sølbeck Rasmussen:** Conceptualization, Methodology, Supervision, Project administration, Writing – review and editing, Funding acquisition.

Ania Stenberg Mortensen and Camilla Frost Holm contributed equally to the experimental work. Henriette Lyng Røder and Torben Sølbeck Rasmussen contributed equally to the conceptual development and supervision of the study and to the writing and revision of the manuscript.

## Funding

This work was supported by the Villum Foundation through a grant awarded to H.L.R. (grant no. 34434) and by the Independent Research Fund Denmark through a grant awarded to T.S.R. (grant no. 3123-00064B).

## Declaration of competing interest

The authors declare that they have no known competing financial interests or personal relationships that could have appeared to influence the work reported in this paper.

## Acknowledgements

The authors thank the statistical advisory unit, University of Copenhagen, for advice on the statistical design, analyses, and interpretation of the results.

## Notes

### Competing Interest Statement

The authors have declared no competing interest.

### Summary of Updates

Especially methods have been adjusted and clarified. Reframed our claims and terminology to reflect responses to fecal virome enriched fractions rather than proven active phage infection, explicitly noting the lack of virome characterization and infectivity testing. Expanded the statistical analysis by adding a linear mixed-effects model on continuous biofilm biomass data (OD590), supporting our initial categorical findings. Clarified experimental details and quality controls, including negative extraction controls, mock communities, media only controls, donor criteria, and ethical approval.

