## Supplementary file for "Fecal virome-enriched fractions are associated with increased biofilm biomass and distinct bacterial responses in biofilm and planktonic communities in an *in vitro* colon model"

**Contents**

**Supplementary Table S1.** PERMANOVA and PERMDISP analyses of bacterial community composition.

**Supplementary Figure S1.** PCoA plots of donor feces, CoMiniGut communities, and background microbiota.

**Supplementary Figure S2.** Differentially abundant taxa in biofilm-associated communities following virome addition.

**Supplementary** Table S1. *Analysis of differences in bacterial community composition (PERMANOVA) and within-group community dispersion (PERMDISP) of CoMiniGut-fermented samples. Variables tested included virome treatment (No Virome, Virome 1–3), sample type (biofilm or planktonic), machine (CoMiniGut system 1 or 2), year (1 or 2), and donor (1, 2, or 3). R² indicates the proportion of variation in community composition explained by each variable. F denotes the pseudo-F statistic for PERMANOVA and the F statistic for PERMDISP. Significant P-values (< 0.05) are shown in bold.*

| **Variable** | **PERMANOVA R²** | **PERMANOVA F** | **PERMANOVA P** | **PERMDISP F** | **PERMDISP P** |
| --- | --- | --- | --- | --- | --- |
| Virome | 0.024 | 1.57 | **0.029** | 0.09 | 0.76 |
| Sample type | 0.069 | 13.5 | **<0.001** | 27.4 | **<0.001** |
| Virome × Sample type | 0.0089 | 0.58 | 0.99 | 3.86 | **0.003** |
| Machine | 0.0076 | 1.49 | 0.13 | 2.73 | 0.10 |
| Year | 0.10 | 19.8 | **<0.001** | 0.13 | 0.72 |
| Donor | 0.36 | 35.6 | **<0.001** | 6.66 | **0.002** |
| Residuals | 0.44 | – | – | – | – |

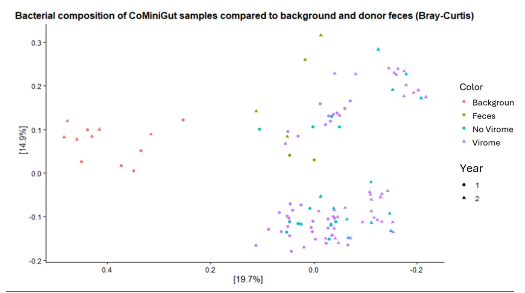

***Supplementary Figure S1.*** *Principal coordinates analysis (PCoA) based on Bray-Curtis dissimilarity showing the bacterial community composition of CoMiniGut background microbiota, original donor feces, and CoMiniGut-fermented samples from the No Virome and Virome treatments. Colors indicate sample category (CoMiniGut background microbiota, donor feces, No Virome, and Virome), and symbols indicate experimental year.*

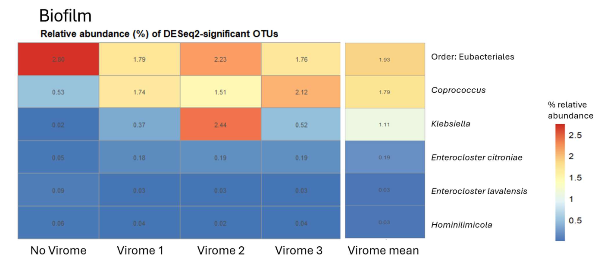
**Supplementary Figure S2.** *Heatmap showing the relative abundance of dominant bacterial taxa that were significantly differentially abundant between No Virome and Virome treatments within biofilm-associated communities (DESeq2; two-sided Wald test; FDR-adjusted P < 0.05; absolute log2 fold change > 0.5). When taxa could not be assigned to genus level, the nearest available taxonomic rank is shown.*

*.*
